# Daily intermittent fasting is an effective multiscale treatment in preclinical models of absence epilepsy

**DOI:** 10.1101/2025.01.21.634091

**Authors:** Coline Rulhe, Steeve Thirard, Juri Aparicio Arias, Leila Makrini, Florence Fauvelle, Emmanuelle Beyne, Alexandra Dauvé, Tristan Bouschet, Emmanuel Valjent, Federica Bertaso

**Affiliations:** Department of Neuroscience, Institut de Génomique Fonctionnelle, Univ Montpellier, CNRS, INSERM, Montpellier, France; Université Grenoble-Alpes, INSERM, CNRS, CHU Grenoble Alpes IRMaGe, Grenoble, France; Montpellier GenomiX (MGX), Université de Montpellier, CNRS, INSERM, Montpellier, France; Current address: INM, Univ Montpellier, INSERM, Montpellier, France

## Abstract

Absence epilepsy (AE) is characterized by brief but frequent seizures with loss of consciousness. Existing treatments have heavy side effects, are only partially effective and do not address the comorbidities, including cognitive and social deficits. A tripartite link between seizures, cognitive deficits and diet has been established. We focused on intermittent fasting (IF), a regime where daily periods of fasting alternate with periods of food intake, with no restrictions in the type or quantity of food consumed. To date, the effects of IF on infantile epilepsy have not been addressed. We evaluated the therapeutic potential of a daily, one-month protocol of IF on three established mouse models of AE: the Grm7^AAA^ KI mouse, the Scn2a haploinsufficiency mouse and the pharmacologically-induced AY-9944 mouse model. We show a reduction of the seizure frequency in all models, as well as an improvement of the sociability deficits observed in two of the models, with no adverse effects. Focusing on the Grm7^AAA^ KI model, we performed RNA sequencing in a one of the key brain areas of the absence seizure circuit, the thalamus. We detected a deregulation of genes involved in vascularization associated with the development of malformed blood vessels in epileptic mice. Along with its anti-seizure effects, IF was able to counteract both abnormal gene expression and vessel morphology. This study demonstrates for the first time the positive effects of IF on AE and could facilitate the implementation of the diet in clinical trials.

## INTRODUCTION

Throughout evolution, food scarcity has driven the development of brain and body functions. While diet, exercise and proper sleep help maintain a healthy lifestyle, overnutrition and a sedentary behavior are a major risk for a variety of pathologies. Intermittent Fasting (IF) is the dietary practice of cycling between windows of eating and abstaining from calorie intake. Different protocols of IF have been developed over the years: Time-Restricted Feeding (TRF) consists in fasting daily for a period, typically 12 to 18 consecutive hours, while Alternate Day Fasting and Periodic Fasting introduce full days of fast throughout the week. Recently, IF has gained popularity, being a simple and cost-free regime, more easily implemented and tolerated compared to other diets. IF has been thoroughly investigated in the context of energy balance and is an efficient weight-loss method, positively regulating blood glucose and insulin levels, and modulating hormone signaling (1). Recent studies have demonstrated a beneficial role of IF on aging in a variety of organisms, ranging from flies (2), mice and rats (3, 4), to healthy humans who display improved health markers (5). In addition, beneficial effects of fasting have been reported in cancer patients and models, notably increased survival (6), delayed tumor progression (7) and reduced adverse effects of chemotherapy (8). For neurodegenerative disorders such as Alzheimer’s, Parkinson’s, Huntington’s disease and multiple sclerosis, IF exerts neuroprotective effects in animal models and improved cognitive functions in patients (9).

Amongst neurological disorders, epilepsy, which affects 50 to 65 million people worldwide (10, 11), is ranked the 7^th^ most impactful in terms of disability-adjusted life-years (12). This central position is emphasized by the World Health Organization’s global plan of action, which makes the access to treatments for epilepsy and other neurological disease a priority (13).

Epilepsy is characterized by the chronic occurrence of spontaneous seizures arising from abnormal neuronal hyperactivity. They can manifest in several ways depending on the affected brain region. Absence Epilepsy (AE) is a common, idiopathic and non-convulsive form of epilepsy affecting children. In patients, mutations leading to the development of AE have been identified, including CACNA1H, GABRA1, GABRG2 and SLC2A1, which account for 10% of cases in children younger than 4 years old at the onset of the pathology. AE is characterized by generalized seizures lasting a few seconds, and by the electrographic signature of spike and wave discharges (SWD) on the electroencephalogram (EEG). During AE seizures, which can occur up to a few hundred times a day, patients experience short lapses of consciousness. It is estimated that 60% of patients are affected by comorbid neuropsychiatric disorders. The prevalence of attention deficits in AE patients ranges from 30 to 60% (14, 15). Patients can also exhibit cognitive disorders and psychosocial difficulties such as anxiety and social interaction problems. Anti-epileptic drugs (AED) for the treatment of absence seizures traditionally involve the use of ethosuximide, a sodium channel blocker, as a first-line treatment. In case of failure, ethosuximide can be used in combination with either valproic acid or lamotrigine. Deplorably, studies report that between 10 and 30% of AE patients are pharmacoresistant and do not respond to AED (16). In addition, the use of drug treatments can be associated with adverse effects, such as fatigue or digestive issues, and may induce or exacerbate comorbid cognitive and psychological conditions (17). Psychiatric comorbidities may persist in patients even after their seizures have been fully controlled. Importantly, while AE seizures remit in about 50 to 85% of patients when they age (18), the comorbidities persist into adulthood. In other patients, the pathology may evolve into other types of idiopathic epilepsy.

Absence epilepsy affects children at a critical period of their development; it is therefore crucial to explore new therapeutic approaches capable of controlling seizure while limiting their side effects. Historically, diets designed to mimic the effects of fasting have been used to treat epilepsy well before the development of AED, the first reports dating back to Hippocrates and his "Sacred Disease" seminal text (19). In particular, the low carbohydrates, high-fat ketogenic diet (KD) was developed in the 1920’s to mimic the metabolic switch induced by starvation. To date, KD remains a mainstay treatment for refractory epilepsy, especially in children (20). However, adverse effects, restrictiveness and reported unpalatability result in low implementation and long-term compliance to KD. While short-term side effects, such as nausea, vomiting or ketosis, are debilitating for patients, durable adverse effects can severely affect their health, as hypercholesterolemia, liver and kidney malfunctions or nutrient deficiencies are reported (21).

Conversely, studies addressing the use of IF as a treatment for epilepsy are sparse and mainly centered on tonico-clonic or temporal lobe epilepsy animal models. For example, in a pilocarpine-induced seizure rat paradigm, a strict TRF based on a 2-hour daily access to food shows an anticonvulsant effect (22), accompanied by a substantial weight loss, while a few preclinical studies report variable effects of alternate day IF (food provided every other day) depending on the model under study (intraperitoneal injection of kainic acid *vs.* 6 Hz threshold test, (23, 24)). However, research on the use of IF on AE has been elusive, with one prospective study reporting positive effects of religious IF on absence seizures (25). Furthermore, the effects of IF on the neuropsychiatric comorbidities found in AE have not been investigated.

In the present study, we combined electroencephalography, behavior analysis, RNA sequencing and histology to decipher the effects of IF on AE in three preclinical mouse models of the AE: the genetic Grm7^AAA^ knock-in (KI) and the Scn2a haploinsufficiency mouse models (26–28), as well as the pharmacological AY-9944 model (29). All three models exhibit absence-like seizures with electrographic SWD, but display complex and distinct neuropsychiatric-like comorbidities. We report that a 1-month IF regime reduced the seizure occurrence in all three models while improving behavioral comorbidities without noticeable side effects. In addition, we found that Grm7^AAA^ KI mice exhibit transcriptomic alterations associated with an abnormal vascular structure and that IF normalizes these patterns at a transcriptional and functional level. Our results support the potential of IF as an effective treatment to approach the pathology in its entirety, addressing the neurological, psychological and social burdens of AE.

## RESULTS

### IF reduces AE seizure occurrence

To assess the effects of IF on *typical* and *atypical* absence seizures and the associated comorbidities described in humans (30), we selected 3 well-characterized mouse models covering a broad spectrum of AE features (Fig. 1A). The Grm7^AAA^ KI/KI mouse model (Grm7^AAA^ KI for simplicity) displays *typical* absence-like seizures with ≍ 6 Hz SWD accompanied by behavioral arrest. Mild comorbidities have been reported so far, including some working memory deficits (28). The Scn2a^+/-^ and AY-9944 (AY for simplicity) are both models of *atypical* absence seizures, with complex behavioral alterations including age-dependent autistic traits (31–34). Control and epileptic mice of the 3 models were randomly assigned to either AL (*Ad Libitum*, unrestricted access to food) or IF (time-restricted access to food for 8 h during the animal’s active phase) diet (Fig. 1B). Both daily food intake and body weight were slightly reduced on day 1 of diet implementation, as the animals adjust their feeding to the food availability schedule. No significant differences were observed between the AL and IF regimes over the following days (Fig. 1 C, D and Fig. S1). We confirmed that the fast-induced gene *Hmgcs2* is upregulated by IF intervention (Fig. 1E). *Hmgcs2* codes for a mitochondrial enzyme that catalyzes the first reaction of ketogenesis, in which ketone bodies are synthetized from lipids to be used as energy during times of carbohydrate deprivation, such as fasting. In the Grm7^AAA^ KI and Scn2a^+/-^ models, *Hmgcs2* was significantly increased in fasted conditions regardless of the genotype. A similar trend was observed in the AY model, although it was not significant. Surprisingly, in NaCl-treated controls, *Hmgcs2* levels were not affected by IF.

**Figure 1:**
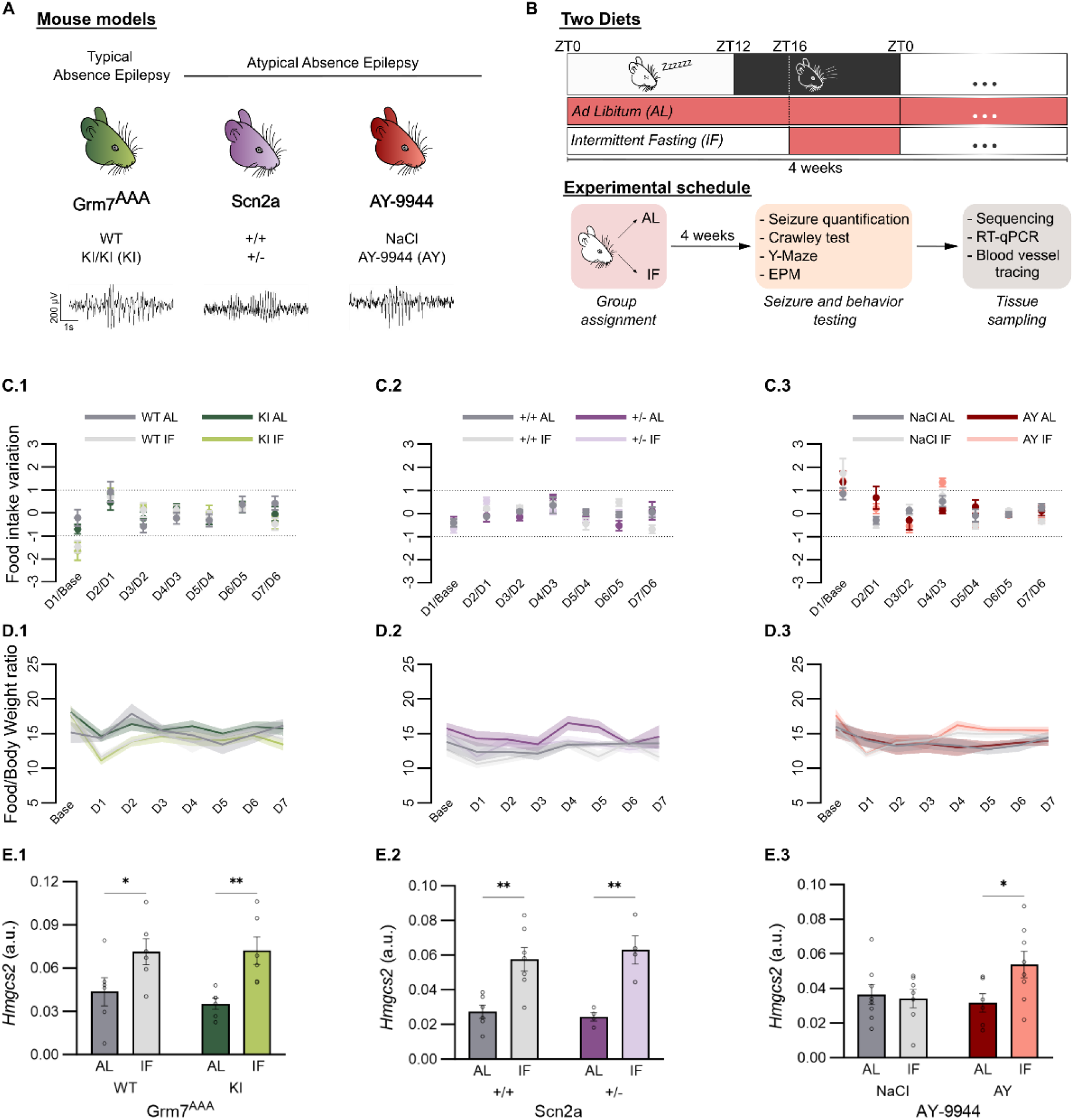
Daily intermittent fasting implementation. A) Mouse models used in the study and example traces of EEG recorded spike-and-wave discharges representative of the models. B) Experimental schedule. C1-3) Food intake variation over the first 7 days of IF. D1-3) Food-to-body weight ratio over the same period of time. E1-3) Relative quantification of *Hmgcs2* mRNA quantification in animals undergoing AL or IF. * p<0.5; ** p<0.01; *** p<0.001, Two-way ANOVA followed by Fisher’s LSD test.

Electroencephalographic (EEG) activity and behaviors were evaluated in AL and IF groups. No abnormal EEG patterns were observed in control animals (Grm7^AAA^ WT, Scn2a^+/+^ and NaCl mice; data not shown) regardless of the diet group. IF *per se* did not induce abnormal EEG signals and no side effects were observed on the animals’ general behavior. Consistent with previous studies, epileptic mice of all 3 models displayed absence-like seizures in the form of spike-and-wave discharges (SWD) with characteristic frequency profiles (Fig. 2A). Grm7^AAA^ KI and Scn2A^+/-^ mice on the IF regime for 1 month displayed a lower seizure rate compared with AL-fed mice (Fig. 2B). The AY model displayed a similar trend, despite not reaching statistical significance. Mice were followed longitudinally before and after the diet to assess the intrinsic evolution of seizure occurrence. This revealed a general worsening of seizures frequency over time for all the AL-fed groups, and a significant reduction for two of the epilepsy models under IF regime, and a tendency for the Scn2a^+/-^ model (Fig. 2C). In summary, IF prevented the worsening in seizure occurrence associated with aging and alleviated the seizure burden in three preclinical models of absence epilepsy.

**Figure 2:**
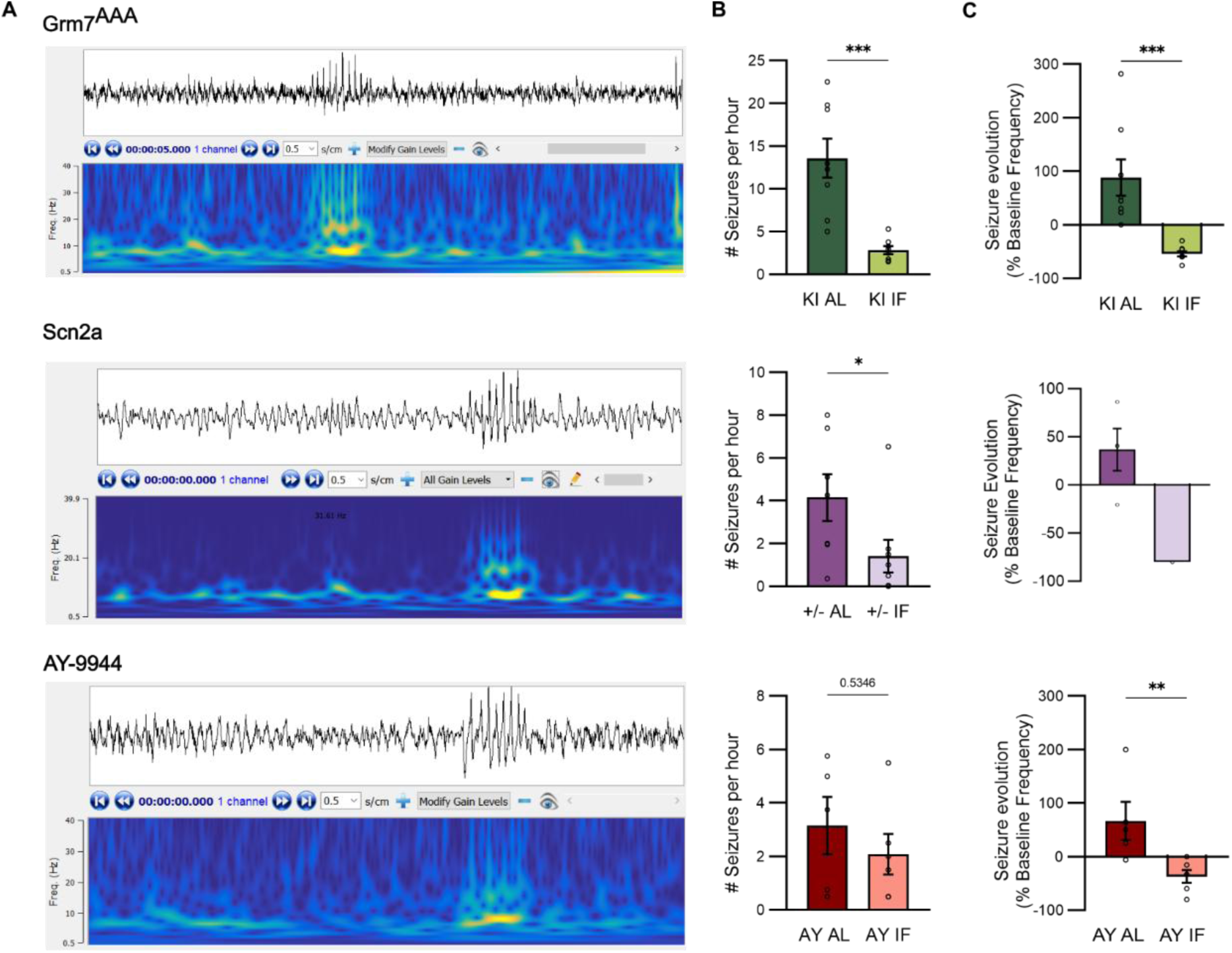
Absence epileptic seizure load decreases after 1-month IF. A) Example EEG recording and wavelet transform analysis for Grm7^AAA^ KI, Scn2a+/- and AY-9944 mice. B) Comparison of seizure load in AL-fed and IF-fed animals of each strain, after 1 month of regime. C) Longitudinal analysis before and after regime implementation, showing the evolution of seizure occurrence. * p<0.5; ** p<0.01; *** p<0.001, Mann-Whitney test.

### IF reverses social interaction deficits with no additional side effects

We then evaluated the impact of IF on behavioral comorbidities associated with AE. Distinct behavioral impairments have been previously reported in the 3 models, including social interaction deficits in the AY model, and working memory deficits in the Scn2a^+/-^ and Grm7^AAA^ KI models. We chose to assess social behavior (Fig. 3A), anxiety-like behavior (Fig. 3E) and working memory (Fig. 3I) during the animals’ active phase (dark phase). In addition, the tests were performed at least 4 h after the IF group was given access to food, to limit exploratory food-seeking behavioral bias. Control animals of both the AL and IF groups were included in the comparisons, as fasting *per se* might represent a stressful challenge for the mice.

**Figure 3:**
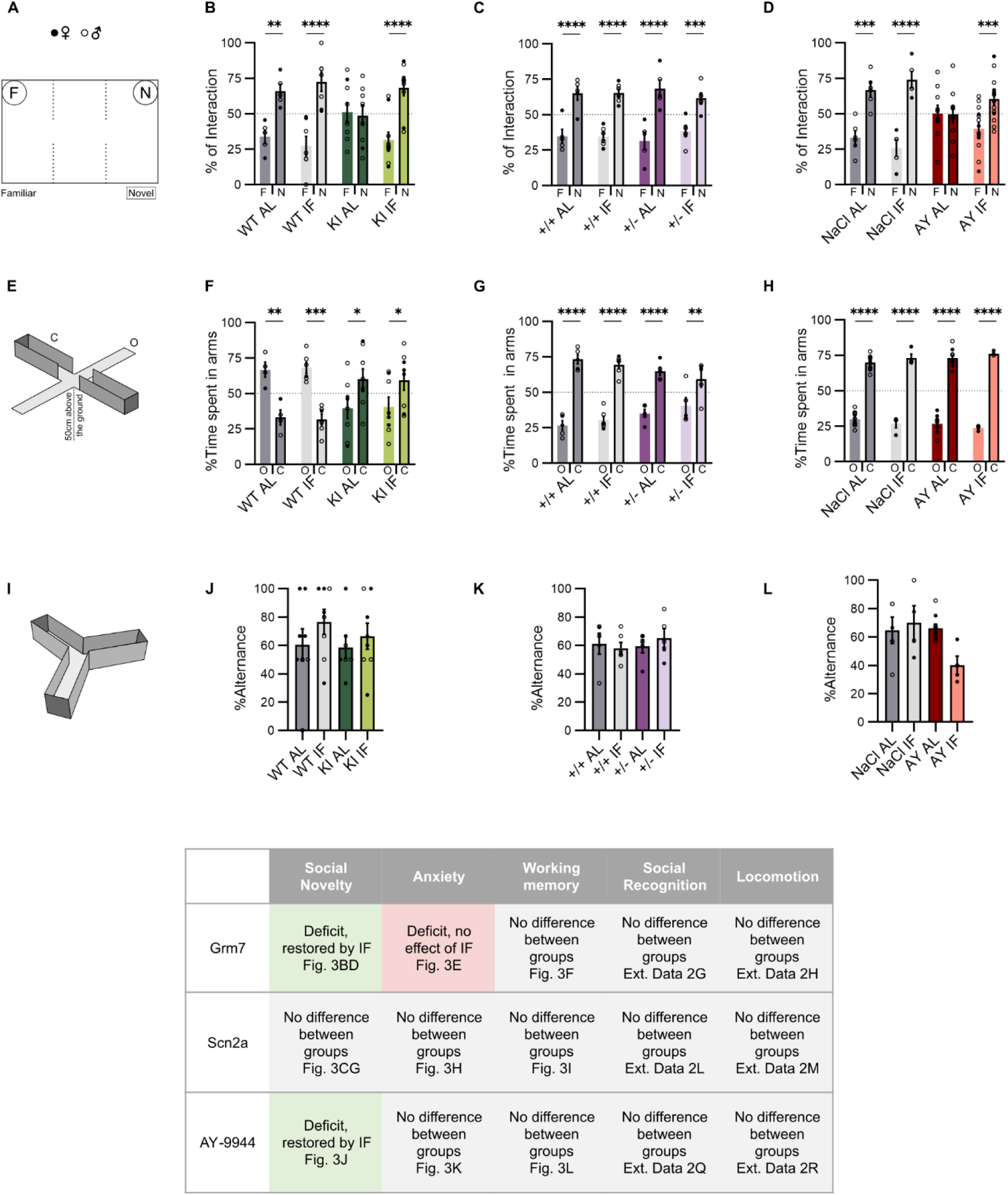
Behavioral comorbidities are improved or unaffected by intermittent fasting. A) Crawley’s three-chamber social interaction test arena. One cage contains a familiar, previously explored mouse (F), the other contains a novel, non-encountered mouse (N). B, C, D) Percentage of interaction with the familiar or novel mouse for each mouse strain and regime. E) Elevated plus maze apparatus with open (O) and closed (C) arms. F, G, H) Percentage of time spent in open (O) and closed (C) arms for the same mouse strains and regimes. I) Y maze apparatus. J, K, L) Quantification of correct alternation between the three arms of the maze for the same groups. * p<0.5; ** p<0.01; *** p<0.001, Two-way ANOVA followed by Tukey’s multiple comparisons test.

In the Crawley’s 3-chamber social novelty test, we show that control animals, regardless of their regime (AL or IF), spent more time interacting with a novel mouse than a familiar one, reflecting a preference for social novelty (Fig. 3A). In contrast, Grm7^AAA^ KI mice showed no preference, revealing a deficit in sociability (50.2% of the time spent interacting with the novel stimuli *vs.* 66.8% for WT). Strikingly, this lack of preference for social novelty was reversed by IF, with interaction patterns restored to control levels (Fig. 3B). Importantly, the total time spent interacting with either stimulus was not affected by the group or the intervention (Fig. S2), excluding a reduction in exploratory activity. Nose-to-nose contact duration was also unaffected in all groups. As previously reported, no social interaction deficits were observed in the Scn2A^+/-^ model, and this was unaffected by IF (Fig. 3C). AY mice displayed the reported lack of social novelty preference as opposed to controls (32), and preference for social novelty was restored to control levels by IF (Fig. 3D).

Next, to assess anxiety-like traits, the mice were tested in an elevated plus maze (Fig. 3E). Epileptic Grm7^AAA^ KI animals exhibited a preference towards the closed arm (where they spent 66.4% of the time, Fig. 3F), reflecting an anxious-like phenotype, as opposed to controls who preferred the open arm (where they spent 60.3% of the time). This behavior was unaffected by the IF intervention in either genotype. Conversely, both Scn2a^+/-^ and AY animals showed a marked preference for exploring the open arms, and no differences were found regardless of group or treatment (Fig. 3G, H).

Finally, we evaluated spatial working memory with a Y-maze (Fig. 3I). The test revealed no alteration in working memory no matter the model tested, and the results were unaffected by IF intervention (Fig. 3J, K, L).

Taken together, these results show that IF can successfully reverse the social interaction deficits exhibited by the Grm7 ^AAA^ KI and AY models (summarized in Fig. 3, bottom table).

Analysis of the transcriptome reveals abnormal synaptic transmission and angiogenesis in the thalami of epileptic Grm7^AAA^ KI mice Because of the nature of the intervention, the effects of IF have been mainly described in the periphery, notably in the liver because of its key regulatory role in metabolism (1). Conversely, while the mechanisms of IF in the healthy brain have been well characterized, a limited number of studies have addressed its effects in the context of epilepsy. A plethora of electrophysiological studies have highlighted the fundamental involvement of the thalamocortical circuit in the generation of absence seizures. To decipher the mechanisms underlying the positive effects of IF on seizures and comorbidities we observed, we conducted an unbiased transcriptome analysis of the thalami to identify differentially expressed genes (DEGs) in our genetic model of typical absence seizures, Grm7^AAA^ KI, and its controls. In the thalami of Grm7^AAA^ KI compared to WT mice (both fed AL) we identified 232 differentially expressed genes (DEGs), including 168 up- and 64 downregulated genes (Fig. 4A-C). The top (Log2(FC)) up- and down-regulated were *Slc16a8* (coding for the monocarboxylate transporter 3) and *Ccl21b* (coding for C-C Motif Chemokine Ligand 21) respectively. A gene ontology (GO) analysis was performed to identify the biological processes associated with the DEGs. The upregulated genes were associated with changes in angiogenesis (5 out of the top 20 GO terms), morphogenesis/BMP signaling (9 out of 20 top G), and motility (6 out of 20) (Fig. 4D, E). On the other hand, GO analysis on the downregulated genes yielded terms associated with synaptic signaling (Fig. 4F, G). Examples of genes found in the upregulated GO term “angiogenesis” and downregulated “synaptic signaling” are depicted in Fig. 4C. Thus, transcriptomic analyses revealed that the AE observed in Grm7^AAA^ KI mice is associated with a defect in synaptic transmission and abnormal angiogenesis.

**Figure 4:**
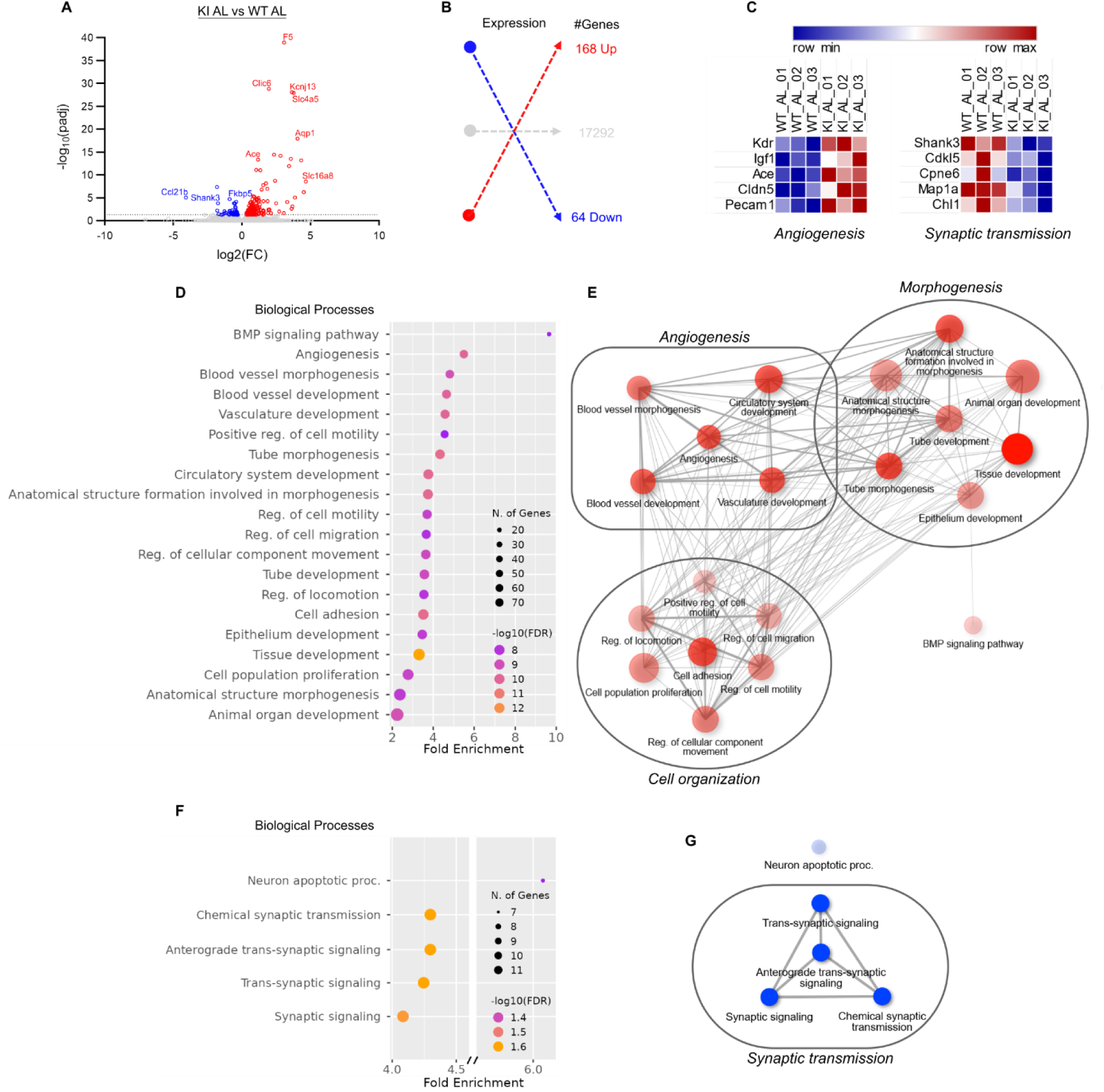
Angiogenesis and synaptic transmission are altered in the thalami of Grm7^AAA^ KI mice. A) Volcano plot illustrating the distribution of gene level in RNA-seq data from Grm7^AAA^ KI and WT controls. Genes that pass the thresholds for FDR and Log Fold Change (identified using DESeq2, p-value<0.05) are colored in red (upregulated genes) and blue (downregulated genes). B) Number of DEGs up- or downregulated, from panel A. C) Heat-maps showing the expression of five genes associated with angiogenesis (left panel) or with synaptic transmission (right panel) in the three WT and three Grm7^AAA^ KI samples. D-F) GO analysis of the biological processes enriched in genes <u>upregulated</u> (D) and <u>downregulated</u> (F) in Grm7^AAA^ KI vs WT animals, and illustrated in the corresponding networks (E, G).

Of note, preliminary data obtained from sequencing thalamic tissue in the Scn2a model show that a very small number of genes are up- or downregulated in epileptic mice compared with controls (data not shown). These results suggest that, despite the central role of Scn2A in determining the pathological phenotype of mice and patients, its effects do not seem to stem from major changes in gene expression in the thalamus.

IF has no major effect on WT mice but partially rescues abnormal transcription patterns in epileptic Grm7^AAA^ KI mice First, we evaluated the effects of IF in Grm7^AAA^ WT mice. We identified a limited set of genes whose expression was significantly modified by the regime: 15 upregulated genes, for which GO did not yield any specific processes, and 11 downregulated genes associated with changes in synaptic transmission (Fig. 5A, B, Fig S3 and Fig. S4). Thus, IF *per se* did not seem to have a major impact on gene expression in the thalamus, consistent with its absence of effect on EEG and behavior.

**Figure 5:**
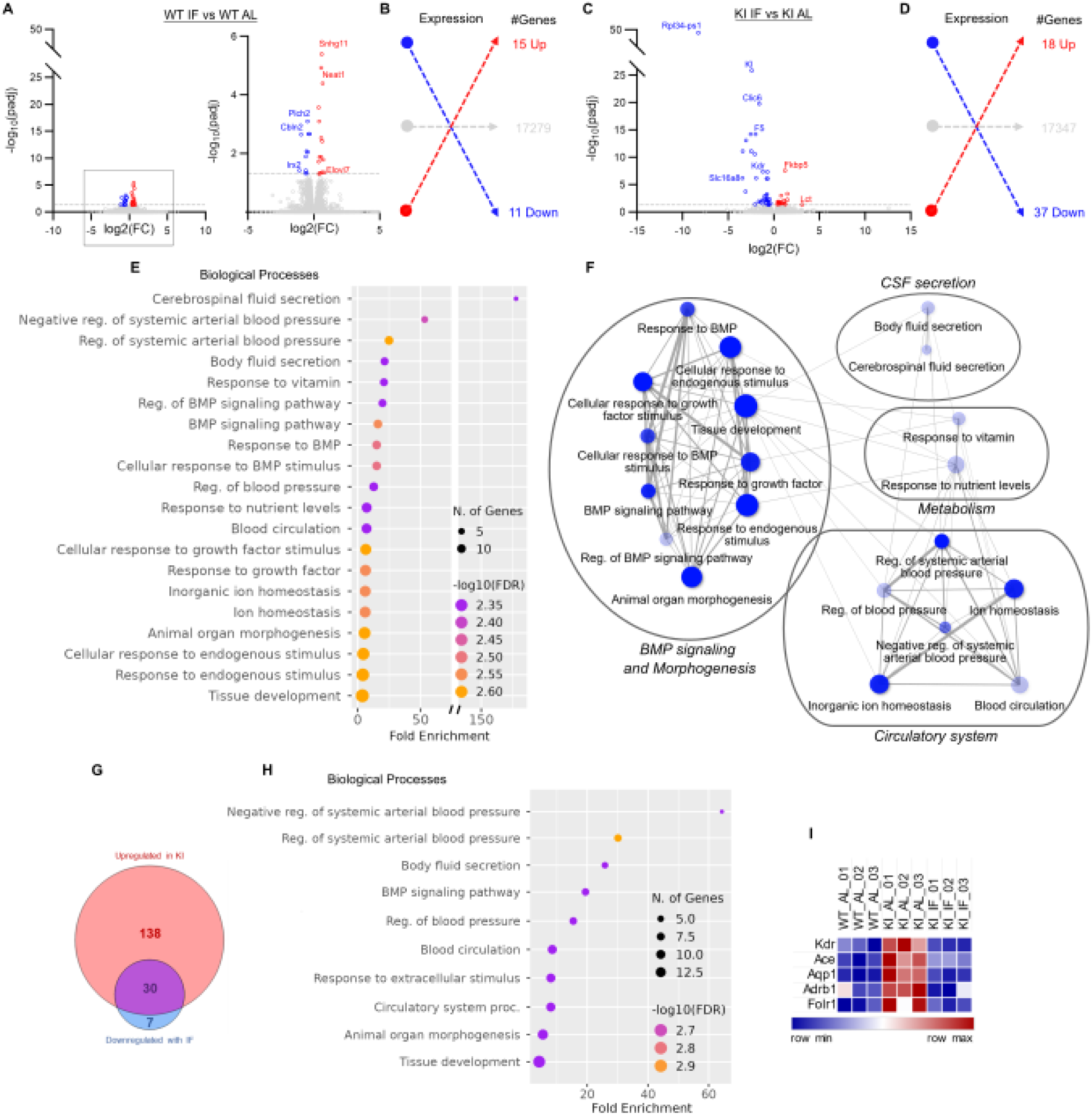
Intermittent fasting reverses thalamic gene expression alterations in Grm7^AAA^ KI mice. A) Volcano plot of RNA-seq data distribution in WT mice AL or IF. Upregulated and downregulated DEGs (DESeq2, p-value<0.05) are colored in red and blue respectively. B) Number of DEGs up- or downregulated, from panel A. C) Volcano plot of RNA-seq data distribution in Grm7^AAA^ KI animals in AL vs IF. D) Number of DEGs up- or downregulated, from panel C. E) GO analysis revealing biological processes associated with DEGs <u>downregulated</u> by IF in Grm7^AAA^ KI animals, and (F) the corresponding networks. G) Venn diagram of the numbers of overlapping genes between KI and. IF conditions. H) GO analysis for the 30 genes <u>upregulated</u> by IF in Grm7^AAA^ KI animals I) Heat-map of selected DEGs found to be upregulated in epileptic Grm7^AAA^ KI and normalized by IF.

We then evaluated the effects of IF in Grm7^AAA^ epileptic animals. In total, we identified 55 DEGs, 18 of them being upregulated and 37 downregulated by IF compared to AL (Fig. 5C, D). The top (Log2(FC)) up- and downregulated were *Lct* (coding for lactase) and *Slc16a8* respectively. While GO analysis on the upregulated terms did not yield any specific results, analysis of the downregulated genes generates terms previously encountered: blood circulation (4 out of the 19 top terms), BMP signaling (4/19) and morphogenesis (4/19, Fig. 5E, F). Interestingly, these terms mirror those associated with upregulated genes in epileptic conditions.

A direct comparison of the genes upregulated in epileptic animals and downregulated after fasting showed that 30 genes were in common: this reveals that 81% of the downregulated genes normalize the biological effects of the altered transcription observed in Grm7^AAA^ KI AL-fed mice (Fig. 5G). GO analysis conducted on the overlapping genes similarly yielded the blood circulation and morphogenesis/BMP signaling pathways, with 6 of the top GO terms in each process (Fig. 5H). The expression of 5 overlapping genes involved in blood circulation, *Kdr*, *Ace*, *Aqp1*, *Adrb1* and *Folr1*, is depicted in Fig. 5I. These data indicate that IF can normalize aberrant gene expression found in epileptic conditions at the transcriptional level.

### IF improves abnormal vasculature in epileptic animals

To determine whether altered gene expression involved in angiogenesis is accompanied by morphological impairments, we investigated the blood vessel architecture in Grm7^AAA^ KI mice. We performed a fluorescence-based tracing experiment in the thalamus to precisely characterize 3 parameters reflecting the complexity of the blood vessels: ramifications, density and total length (Fig. 6A-D). We found that blood vessels in the thalamus of epileptic mice have more ramifications (+48%, 64.4 au *vs.* 43.5 au in controls, Fig. 6E) and a decreased density (-25%, 16.5 *vs.* 22.1 au in controls, Fig. 6F), leading to a smaller total length (-16%, 21.4 *vs.* 25.3 au, Fig. 6H). This is consistent with the transcriptomic results pointing at an abnormal expression of angiogenesis associated genes. Importantly, after 1 month of IF, all 3 of these parameters were rescued to control levels. Lastly, we sought to confirm the expression of *Kdr* through RT-qPCR. After confirming the decrease in *Kdr* expression in the Grm7 model, we assessed it in the Scn2a and AY models. Interestingly, while *Kdr* expression was not altered in epileptic conditions, it was decreased after IF intervention regardless of the group (Fig. 6H and data not shown).

**Figure 6:**
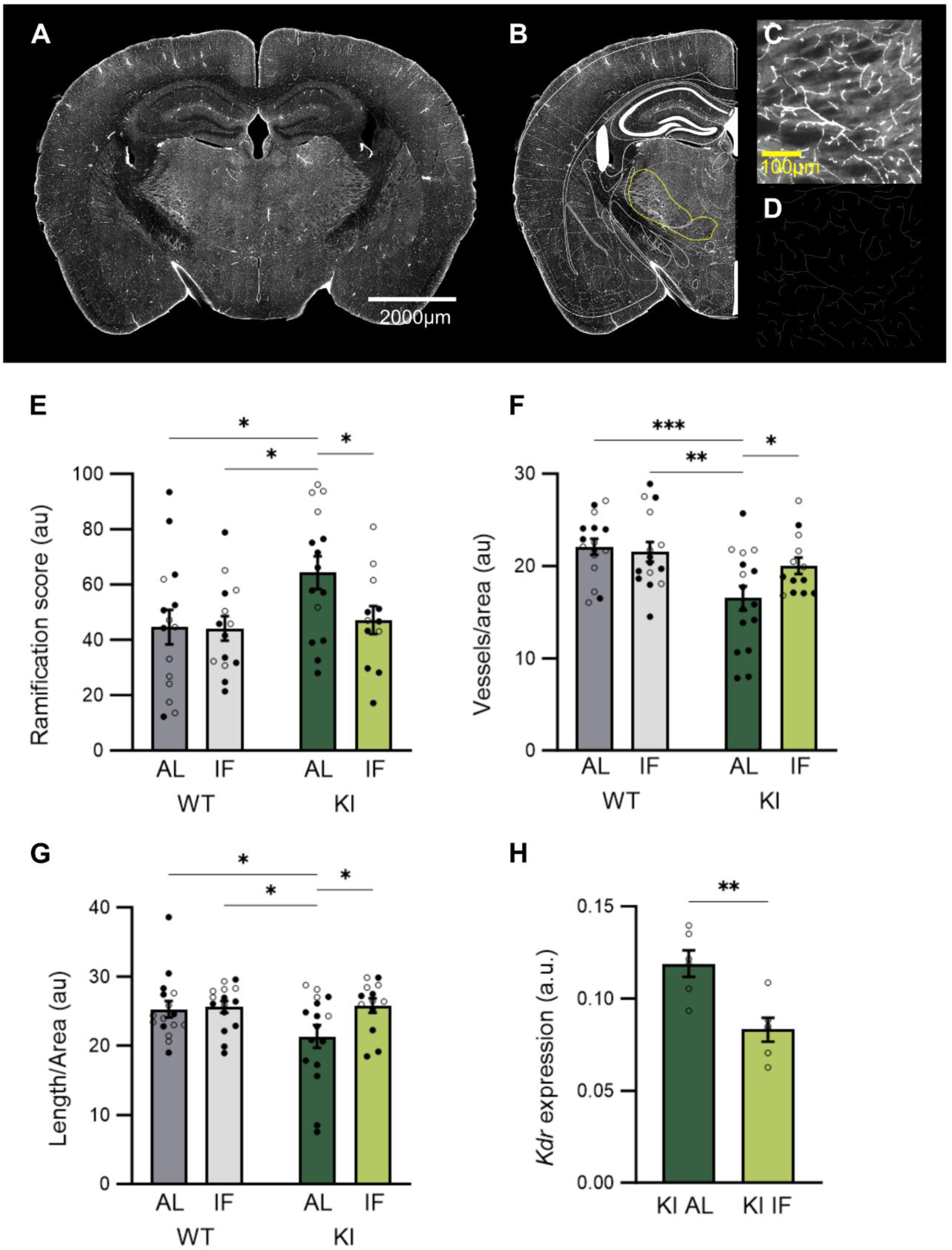
Thalamic vascular architecture differences are normalized by IF in Grm7^AAA^ KI mice. A) Coronal brain slice mosaic image from a FITC-albumin-perfused animal (20X magnification). B) Schematic superposition of brain regions selected for vessel quantification. C) Zoom on thalamic FITC-albumin- labelled vessels. D) Semi-automatic vessel tracing rendering with the *Skeletonize* ImageJ plugin used for vessel parameters quantification. E, F, G) Vascular parameters summary for Grm7^AAA^ WT and KI animals, AL-fed (dark grey, dark green) or IF-fed (light grey, light green). H) Relative quantification of *Kdr* mRNA quantification in Grm7^AAA^ KI animals undergoing AL or IF regimes. * p<0.5; ** p<0.01; *** p<0.001, Two- way ANOVA followed by Fisher’s LSD test.

Taken together, these findings highlight the potential of this regimen as a multi-scale treatment for AE.

## DISCUSSION

Finding treatments for epilepsy is a long-lasting challenge. Despite the efforts to diversify the biological targets for anti-epileptic drug development, a solid 30% of patients remain pharmacoresistant. Contrary to other forms, such as temporal lobe epilepsy, studies investigating non-pharmacological alternatives to treat absence seizures are scarce. Our study shows that IF leads to an improvement in both seizure frequency and behavioral comorbidities in epileptic animals. All three models (Grm7^AAA^ KI, Scn2a^+/-^ and AY-9944) display a worsening in seizure frequency as they age. Our findings support that IF protects against such deterioration, modifying the disease trajectory and leading to a reversal of neurological alterations in three complex preclinical AE models.

Daily fasting induces a cyclic metabolic switch with oscillations of signaling pathways, creating a balance between the induction of cellular stress resistance and neuronal growth and plasticity mechanisms (35). A variety of direct mechanisms have been identified, notably via the KB produced after fatty acid breakdown during the starvation phase. Several studies have described how KB directly modulate neuronal firing rates, by affecting GABA and glutamate signaling, KCNQ channels, or K-ATP and ASIC channels through alterations of ATP/ADP ratio and pH levels respectively (36). Interestingly, we conducted a preliminary metabolomic analysis of thalamic tissue from Grm7^AAA^ KI mice and we observed no significant variations in 18 major brain metabolites, including GABA and glutamine, regardless of genotype or diet (Fig. S5). Nonetheless, our experimental paradigm successfully induces an increase in *Hmgcs2*, coding for the key regulator of ketone bodies (KB) production. Long-term effects of KB include modifications of cell signaling and modulations of gene transcription. While daily KB oscillations may directly modulate excitability, concomitant effects on blood vessels might explain the positive effects of IF in our study. Interestingly, Hartman *et al*. show that KD and a protocol of alternate-day fasting lead to distinctive protective effects against seizures that cannot be solely explained by the processes directly associated with KB (24). Finally, peripheral effects of IF might play a major role in the modulation of the epileptic phenotype. An increasing number of studies hints at a tripartite link between diet interventions, seizures and gut microbiota. For example, a study in mice showed that the anti-seizure effects of the KD were mediated by the gut microbiota (37). The positive effects of IF on AE that we observed are most likely the result of a combination of mechanisms on multiple levels, both at the periphery and in the brain.

Here, we report a vascular alteration in the brain of Grm7^AAA^ KI animals, with the presence of aberrantly ramified blood vessels, and we further show that IF can normalize these alterations. Vascular abnormalities have not been studied in AE, including in paramount models such as GAERS or WAG/Rij rats. However, peri-ictal changes in cerebral hemodynamics have been reported in GAERS rats (38), along with decreased blood flow in cortical capillaries during seizures (39). This study also reports a decrease in cerebral blood flow in the middle cerebral artery of children during absence seizures. Conversely, alterations of the neurovascular unit have been extensively described in other types of epilepsy (40, 41), including vascular abnormalities and excessive angiogenesis in the context of TLE, both in patients and animal models. These studies also relate blood-brain barrier dysfunctions associated with vascular malformations, in particular because of alterations of tight junctions leading to proepileptic leakage.

Our findings highlight that IF can positively remodel blood vessels in the thalamus of epileptic animals, with all measured parameters (ramification, density and length) returning to control levels. These results are consistent with previous studies supporting IF as an efficient modulator of the vasculature in other pathological contexts. Notably, IF was shown to attenuate the vascular damage in a model of vascular cognitive impairment (42). While the literature on the effect of IF on blood vessels remains limited, studies have focused on vasculature remodeling after different types of diet interventions. Interestingly, protocols of a prolonged fasting of 48 h enhanced angiogenesis in an ischemic brain model, improving the long-term motor outcomes in mice. Additionally, multiple studies have looked at the link between ketogenic diet (KD) and vasculature. In a glioma model, KD reduced the tumor microvasculature, which was associated with improved survival in mice (43), while a study on calorie-restricted KD showed that the intervention was able to reduce angiogenesis in a region-dependent manner (44).

Consistent with the morphological alterations observed with the vasculature, we identified a deregulated expression of genes involved in angiogenesis. In epileptic animals, we observed that 46 out of the 168 upregulated genes were associated with the biological process “Vasculature development” (GO:0001944). Several genes of interest have been identified in other types of epilepsy where angiogenesis is altered. Notably, Castañeda-Cabral *et al.* report an alteration of tight junction proteins such as claudin 5, occludin and ZO1, and an alteration in the VEGF pathway. VEGF-mediated angiogenesis has been described as a mainstay of vascular alterations in the context of epilepsy. We report an upregulation of *Kdr* in Grm7^AAA^ KI animals, encoding the VEGF receptor VEGFR2, similar to what is described in both humans and models, including TLE patients (45, 46) and pilocarpine-induced SE in rats (47). VEGF has been described to be released by neurons during seizures and plays multiple roles, including protection against excitotoxicity and increased angiogenesis and hypertrophy of the vessels, priming the environment to meet the increase in metabolic needs. However, the newly formed vasculature has been described as immature, with disorganized and leaky vessels, making the VEGF pathway a double-edged sword. Interestingly, we found that 14/37 genes downregulated after fasting are associated with the biological process “Circulatory system development” (GO:0072359, such as *Kdr*, showing that IF was able to normalize its expression in epileptic conditions. A link between fasting and VEGF was previously demonstrated by Kim *et al.*, where fasting was able to induce a VEGF expression in the white adipose tissue (48). Otherwise, to the best of our knowledge, studies on the dietary interventions such as IF or the KD and their potential effect on the VEGF pathway are sparse. Calorie restriction led to a reduction in VEGF expression in rats implanted with prostate adenocarcinoma cells (49). In humans, a high-fat diet induced an increase in serum VEGF (50), emphasizing the link between diet and the VEGF pathway. Further research is required to study more closely the organization of the vessels and assess whether or not they are mature and functional in Grm7^AAA^ KI animals.

Other biological processes have been identified as dysregulated in epileptic conditions and need further exploration. Notably, we showed that genes associated with the BMP pathway are upregulated in epileptic mice, namely BMP4 and 6. Both genes were reported to be altered in the periphery after fasting (51). The functions of the BMP pathway are broad. The BMP pathway has been shown to play a role in vasculature development, where alterations of its signaling lead to a malformation and fragility of the blood vessels. On the other hand, the BMP pathway has been shown to regulate neuro- and gliogenesis, as well as synapse and spine size, thus modulating neural plasticity. The BMP proteins have been reported to have contradictory roles depending on the pathological context (52, 53). A recent study underlies a direct link between the BMP pathway and epilepsy: BMP2 is upregulated in response to the overactivation of excitatory neocortex neurons and the disruption of the BMP2-Smad pathway leads to the spontaneous development of seizures in mice (54).

Although AE is a generalized form of epilepsy, multiple findings support the fact that typical absence seizures are driven by the thalamocortical network and then propagate to the entirety of the brain. Several studies identify the primary somatosensory cortex as an initiation site for seizures, working together with the thalamus to induce synchronized oscillations within the circuit. In recent years, other regions have been identified as having a driving role in the generation of seizures. The basal ganglia, long believed to merely be a modulator of the thalamocortical circuit (55), has emerged as a driver of absence seizures (56). Notably, the striatum has been demonstrated to promote synchronous oscillatory activity in the AE circuit (56, 57). Moreover, several studies describe an alternative network involved in the generation of atypical seizures, leading to a more severe phenotype. Interestingly, Shin *et al.* showed an alteration of the hippocampus activity in the Scn2A^+/-^ model, linked to comorbid deficits in memory (58). The study of non-canonical regions might be of interest depending on the model and might provide deeper insight into the understanding of neuropsychological comorbidities. In the Grm7^AAA^ KI model, while social interaction deficits were reversed after the intervention, the anxiety-like behavior was unaffected by the diet. Consequently, further work on brain regions involved in anxiety, such as the insular cortex or the amygdala, might give insight into the behavioral findings. To finish, other behaviors remain to be investigated. Mutations in both the *Grm7* and *Scn2a* genes have been identified in ASD patients (59, 60). Juvenile Scn2a^+/-^ mice have been reported to display autistic-like characteristics, such as higher sensory sensitivity to light, repetitive behaviors, and reduced vocalizations (33, 61). In mice, Grm7 deletion, which leads to the development of seizures, is associated with abnormal social behaviors and alterations in associative fear learning (62). Further work is required to assess discreet autistic-like traits and the potential effect of IF in our models.

Taken together, the present findings provide a useful basis for future therapeutic studies. As of now, the scope of clinical studies of IF in the context of AE is limited. One key finding of our study is that IF did not induce differences in food intake or body weight, nor did we observe IF-induced abnormal behaviors in the treated wild-type or epileptic animals. These data support the need for further research to evaluate the tolerability of IF in children since very few studies have addressed the question. Case studies provide encouraging data, with good tolerability and easy implementation being reported in obese pediatric patients (5 to 15 years old, (63)). In addition, religious intermittent fasting does not induce stress nor detrimental effects on cognitive functions in healthy teenagers, despite the adjustments in sleep patterns resulting from the dawn to dusk fast (64).

In conclusion, this study provides insights into the protective effects of IF on AE. We show that IF is a safe, cheap, effective and easy-to-implement approach that improves both the seizure burden and behavioral comorbidities. While further research is required to deepen our understanding of the mechanisms underlying these effects, our work offers a basis for therapeutic studies and might ease the future clinical implementation of IF in children, as a stand-alone treatment or an adjuvant therapy for AE.

## METHODS

### Experimental design

Unless stated otherwise, our study examined male and female animals, and similar findings are reported for both sexes and therefore pooled. Mouse lines were housed and bred in stable conditions at 22°C, with a controlled relative humidity at 60% and a 12h-12h light/dark cycle in individually ventilated units (Emerald IVC, Tecniplast, Italy). All animals had unlimited access to water throughout the experiment and were fed a regular chow (standard diet A04, SAFE Diets, France) *Ad Libitum* (AL) before IF intervention was initiated. Mice were housed in groups (5 animals/cage) before the EEG surgery and were then housed individually to prevent injuries. Individual cages were set into a 24-cages ventilated cabinet where the mice could hear, smell and see their congeners. Their daily light/dark cycle was inverted over 10 days to align with the experimental schedule of IF. At 8 weeks of age, the animals were divided into 2 groups according to a simple randomization method. The AL group kept unlimited access to food, while the IF group were given access to food for 8 hours during their active phase, 4 hours after the start of the dark phase, and then food deprived for the remaining 16 hours (Fig. 1B).

During the 8 hours feeding window, mice were free to consume food with no caloric restriction. This protocol was repeated daily for the 28-day intervention. During the first week, body weight and food consumption were monitored daily. EEG and behavioral analyses were performed before the start and at the end of the IF protocol and were then sacrificed for tissue collection.

### Mouse models

*Grm7^AAA^ KI model*. The generation and characterization of mutant mice were previously described by Zhang *et al.* and Bertaso *et al.* (26, 28). Briefly, this model is a KI mouse mutant for the mGlu7 receptor which displays absence-like seizures. Behavioral comorbidities have been previously identified in this model, notably working memory deficits. Homozygous KI/KI mice were used to study epileptic individuals, while wild-type littermates were used as controls.

*Scn2a^+/-^ model*. The Scn2a^+/-^ is a haploinsufficiency model, with reported cases of mutations in the *SCN2A* gene found in patients (65, 66). Heterozygous mice display absence-like seizures and complex comorbid symptoms such as impaired learning and memory processes. Heterozygous +/- mice were used to study epileptic individuals, while wild-type littermates were used as controls.

*AY model*. The AY-9944 is a pharmacological model inducing chronic seizures in animals. It has been described as a valid AE model, recapitulating both atypical absence-like seizures and social-behavioral comorbidities (32). The induction of AE in animals has been previously described by Cortez *et al.* and Stewart *et al.*, in rats and mice, respectively (29, 31). Briefly, C57/B6J obtained from Janvier were bred to obtain pups which were treated with a subcutaneous injection of AY-9944 (7.5 mg/kg, 10 µl/g) every 6 days from post-natal P2 to P20. Aged-matched pups injected with an equivalent volume of 0.9% NaCl solution were used as controls (mentioned as "NaCl" condition in the text and figures).

### EEG recordings

Young adult male mice (6-7 weeks old) were anesthetized with a mix containing ketamine (Imalgene 500, 100 mg/kg) and xylazine (Rompun 2%, 10 mg/kg) in PBS and placed in a stereotaxic frame using the David Kopf mouse adaptor. Three-channel preamplifier microconnectors (Pinnacle Technology Inc., Lawrence, KS, USA) were surgically implanted onto the mice, and fixed on the skull with dental acrylic cement. Two electromyography wires were inserted into the neck muscle. After surgery, animals were individually housed and left to recover for 10 days before recording. Freely moving animals were put into individual Plexiglas boxes, and their microconnectors were plugged to an EEG preamplifier circuit and the EEG amplifier. They were left to freely explore and get used to the boxes for 2 h, and were then recorded for at least 4 h along with video monitoring of the animals’ behavior. The electrical activity recorded by extradural electrodes was sampled at 400 Hz filtered at 40 Hz, and recorded by a computer equipped with Sirenia® software (Pinnacle Technology Inc.).

### Behavior

All behavioral tests were performed during the active phase, at least 4 hours after the IF group had had access to food to limit food-seeking hyperactivity.

#### Crawley’s 3-chamber test

A 3-chamber arena was used to assess sociability and preference for social novelty in mice. The arena (50x40x60 cm) is composed of 3 chambers divided by Plexiglass walls. A 6 cm opening in each wall allows the mice to freely explore all 3 compartments. Both side chambers contain a round gridded cage to allow contact between the test mouse and the stimuli placed inside the cage. In the 1^st^ trial (habituation phase), the test mouse was placed in the center of the arena containing the 2 empty cages and was left to explore freely for 10 min. The test mouse was then confined in the center chamber while 2 stimuli were placed in the cages: a gender-matched unfamiliar juvenile mouse *vs.* a mouse-shaped object. During the 2^nd^ 10 min trial, sociability was assessed as the test mouse was able to freely interact with either stimulus. During the 3^rd^ and last 10-minute phase, the object was replaced by a new, unfamiliar mouse, and interaction with the stimuli was measured to assess preference for social novelty. The arena was cleaned with 30% ethanol in between sessions. The choice of the interactor and their placement (left or right cage) were alternated for each trial. Trials were recorded and analyzed using the Ethotrack software (Innovation Net, France) to quantify interactions between the test mouse and the stimuli. Mice were considered to be interacting with the stimuli when their nose was directed towards the cage containing the interactor (less than 2 cm away from the grid). In addition, nose-to-nose contact duration between the test mouse and interactor(s) was manually quantified.

#### Elevated Plus Maze

An elevated plus maze was used to assess anxiety-like behaviors in mice. The maze is composed of four 40 cm-long arms, 2 open arms and 2 enclosed arms elevated 50 cm above the ground. At the beginning of the trial, mice were placed in the center, facing the open arm, and were left to freely explore the maze for 5 minutes. The maze was cleaned with 30% ethanol and left to dry completely between sessions. Video recordings were made of each session and the time spent in each arm was manually measured.

#### Y-Maze

A Y-Maze was used to assess spatial working memory in mice. The symmetrical maze is composed of three 35 cm-long arms. At the beginning of the trial mice were placed at the end of one arm and were left to freely explore the maze for 2 min. The maze was cleaned with 30% ethanol and left to dry completely between sessions. Video recordings were made of each session and the number and sequence of entry in each arm was manually assessed.

### RNA-seq and transcriptome analysis

Grm7^AAA^ KI mice and WT mice, under AL or IF regimens, were deeply anesthetized using Euthasol (140 mg/kg). Their thalami were extracted on ice, transferred to cryotubes and immediately immersed in liquid nitrogen. Samples were kept at -80°C until further processing. Total RNAs were extracted using the quick RNA kit (Zymo) and treated with DNAse I on column. RNAs were quantified on a Nanodrop and their profiles were checked on a Bioanalyser (Agilent). Strand-specific RNA sequencing was performed in triplicate (3 WT_AL, 3 WT_IF, 3 Grm7AAA KI_AL, and 3 Grm7AAA KI_IF) by the MGX facility (Montpellier GenomiX) as previously described (67). Differentially expressed genes (DEG) genes were identified using DESeq2 (68) with an FDR (False Discovery Rate) set at 0.5%. Transcriptome analysis was performed on DEGs using the ShinyGo tool (69).

### RT-qPCR

RNAs were retro-transcribed using N6 primers and M-MuLV retro-transcriptase (RT). qPCR was performed in triplicates using per well: 1 ng of retrotranscribed RNA, primer duplexes (0.3 µM) and SYBR Green Mix (Roche) in 384-well plates on a LightCycler480 apparatus (Roche), as described (70). The level of expression of *Kdr* was normalized to the geometric mean of the level of expression of *Gapdh*, *Mrlp32*, and *Tbp*, which were selected as stable housekeeping genes using the geNorm method (71). Primers are listed in Table S1.

### Metabolomic analysis

Thalami extraction was performed in the same way as for the sequencing experiments, frozen in liquid nitrogen and stored at -80°C until analysis. The tissue samples were analyzed without any extraction process using ^1^H HRMAS NMR (High Resolution Magic Angle Spinning Nuclear Magnetic Resonance) at 500 MHz (IRMaGe facility, Grenoble). 18 different metabolites were detected and quantified using the jMRUI software: acetate (ACE), alanine (ALA), ascorbate (ASC), choline (CHO), CR, GABA, glutamate (GLU), glutamine (GLN), glycine (GLY), glycerophosphocholine (GPC), glutathion (GSH), lactate (LAC), myo-inositol (M-INS), N-acetyl aspartate (NAA), phosphorylcholine (PC), phosphoetaloamine (PE), scyllo-inositol (S-INS), and taurine (TAU).

### Blood vessel analysis

Mice were deeply anesthetized using sodic pentobarbital (Euthasol, 140 mg/kg). Mice were then perfused intracardially with an albumin-fluorescein isothiocyanate conjugate solution (FITC-albumin, Sigma Aldrich, France; 25 mg/ml in cold PBS). After 1 minute, the brains were dissected and put in a 4% paraformaldehyde solution for 48 h. 40 µm coronal sections were mounted and imaged on an epifluorescence microscope at 20 magnification. Exposure was kept constant between acquisitions to allow for comparison between the different conditions. Images were processed and analyzed using the open-source NIH ImageJ analysis software. Parameters such as density, length and complexity were assessed using the Skeleton plug-in and normalized to the total area analyzed.

### Statistics

No statistical methods were used to predetermine sample sizes but they were estimated based on previous experiments of similar nature. Mice were arbitrarily assigned to AL or IF groups. With the exception of RNA sequencing, all data are reported as mean±SEM. Significant differences between mean values were determined using the GraphPad Prism 10 software’s statistics routines. Means were considered to be statistically significant if *p*< 0.05. All statistical analyses are summarized in Table 1.

**Table 1.**
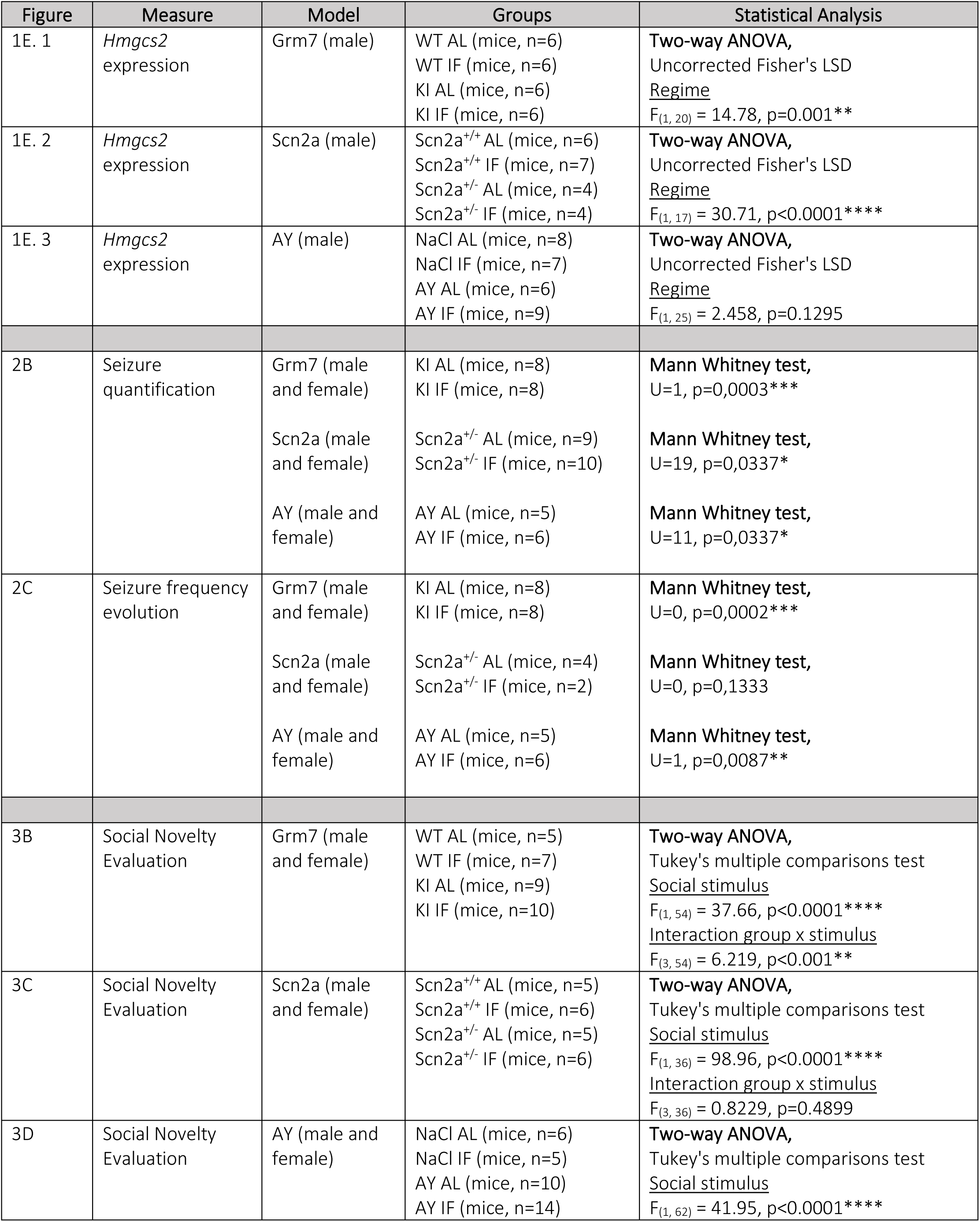
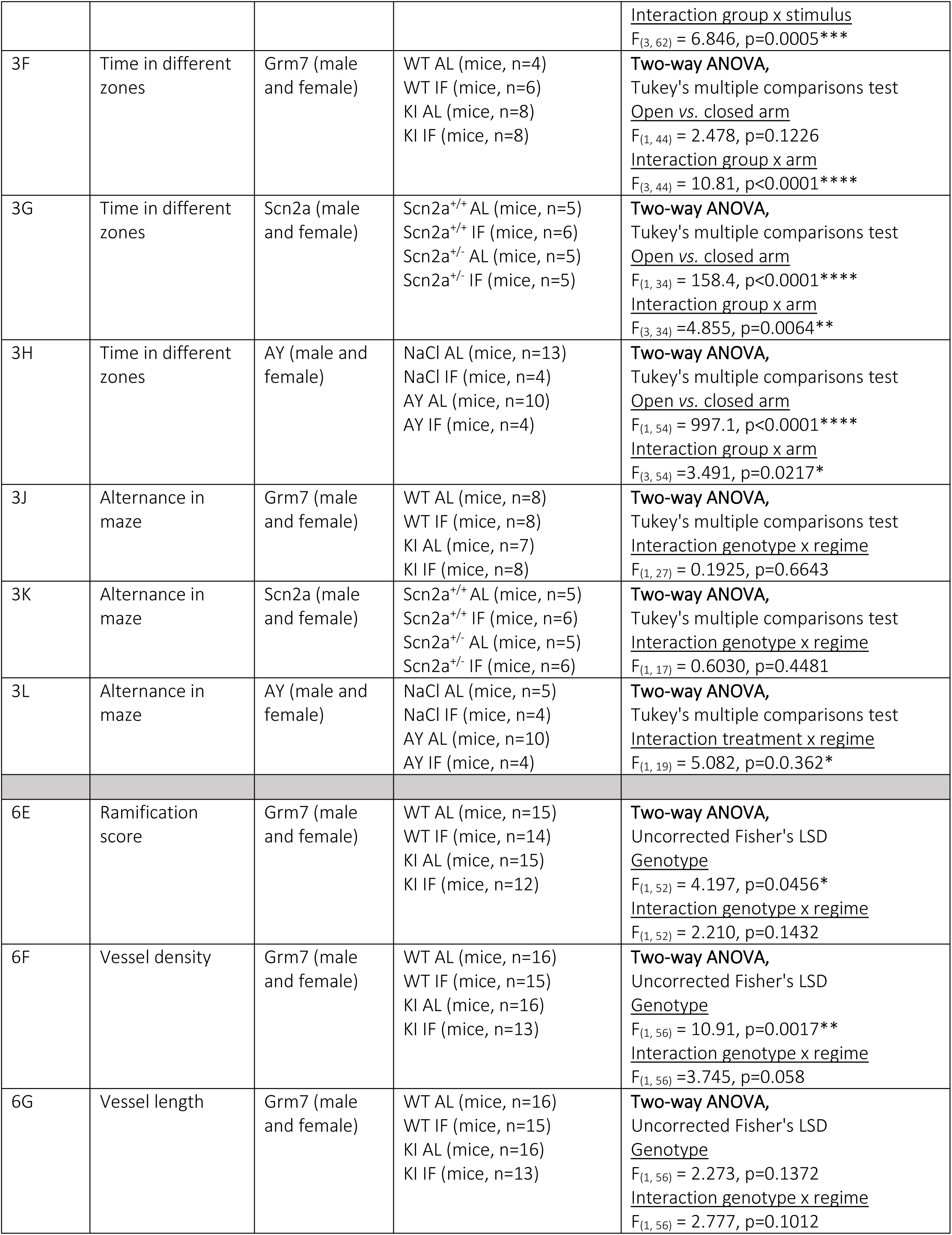
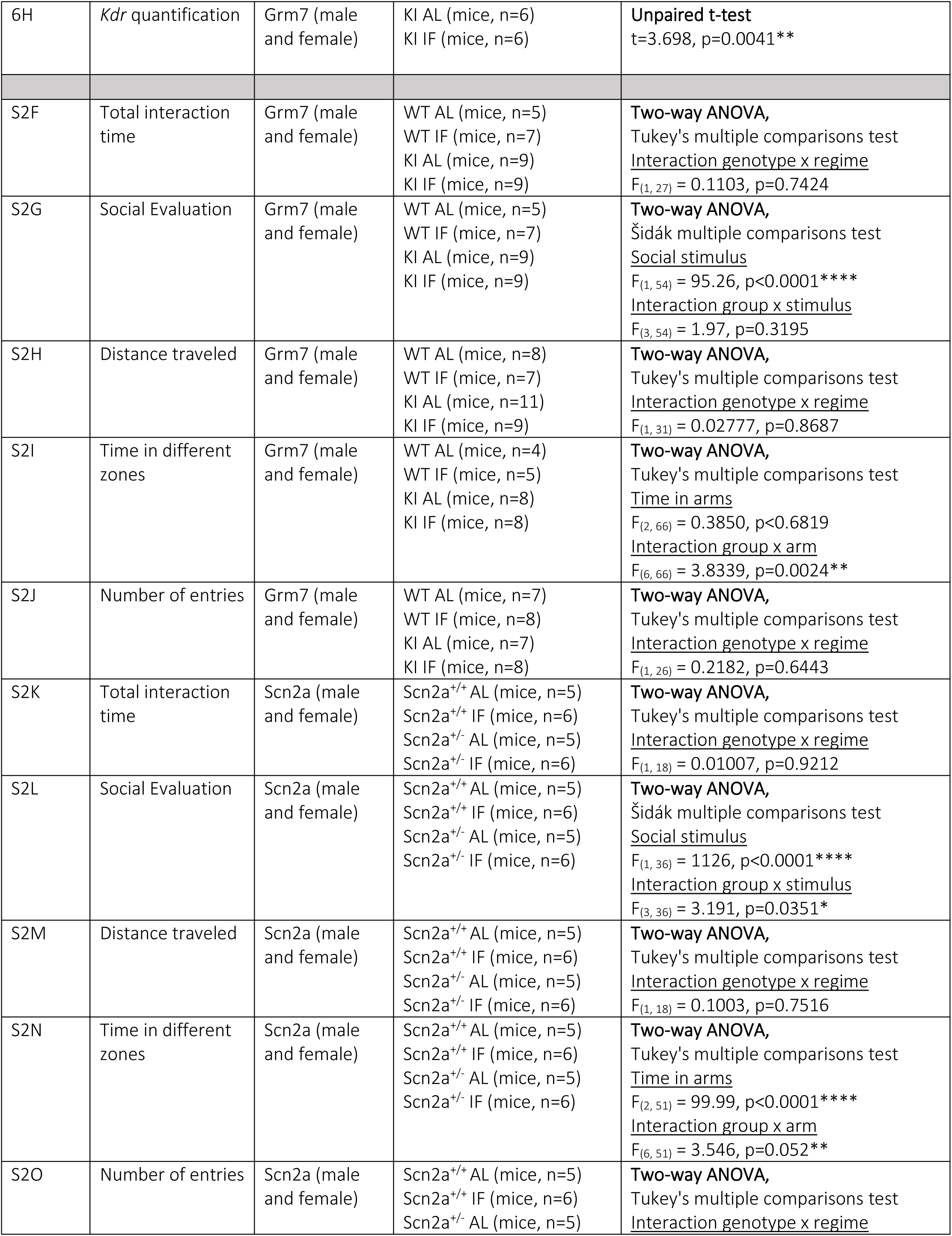
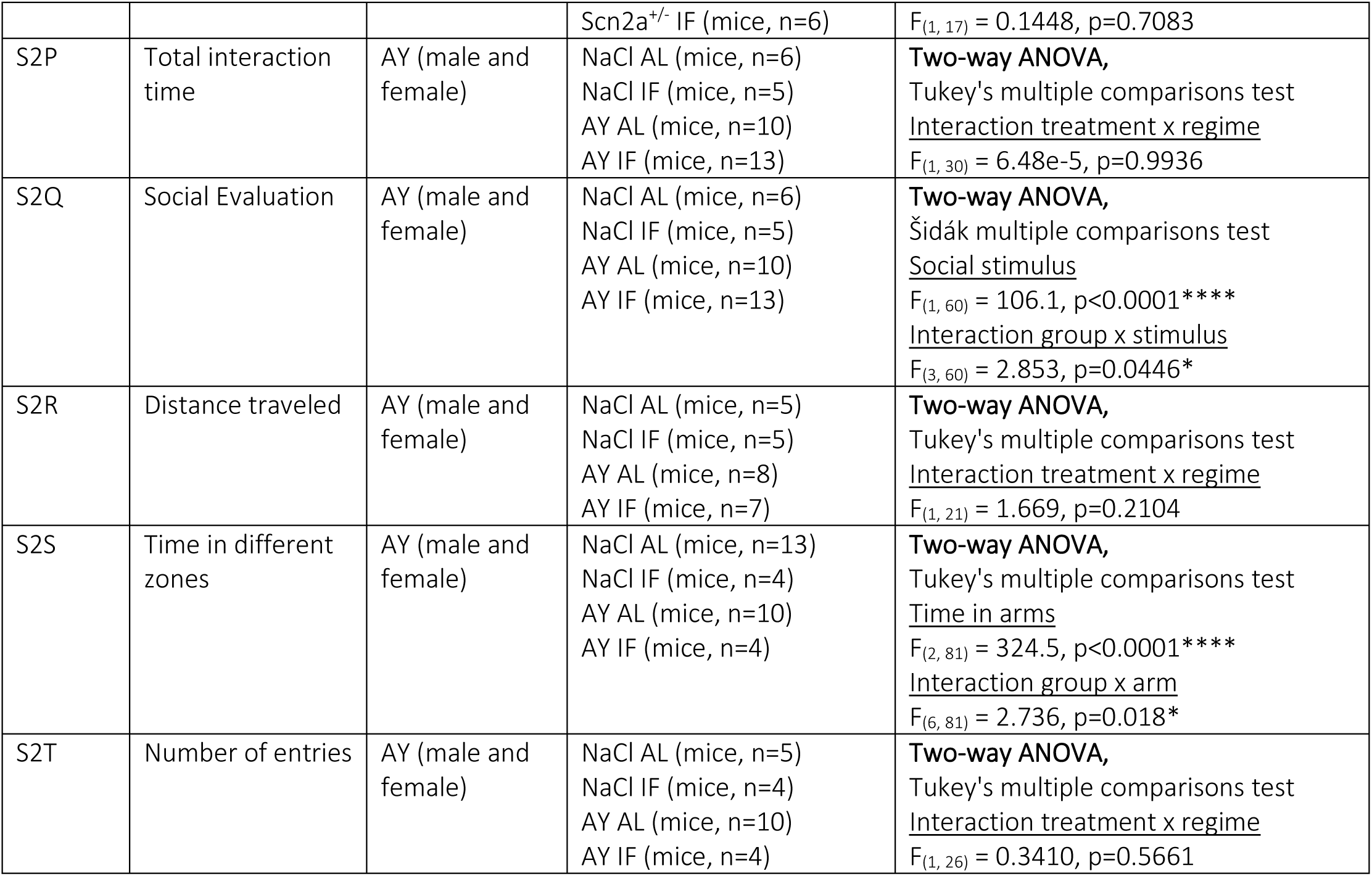
Summary of statistical analyses.

### Study approval

All animal procedures were conducted in accordance with the European Communities Council Directive, approved by the French Ministry for Agriculture (2010/63/EU, authorizations #37118- 2021101411215647 v10 and #37117-202203311212540 v8) and supervised by the IGF institute’s local Animal Welfare Unit.

## AUTHOR CONTRIBUTIONS

Conceptualization was the responsibility of FB and CR. Investigation was carried out by CR, ST, JAA, LM, FF, EB and AD. Formal analysis was performed by CR, FB and TB. Writing of the original draft was carried out by CR. Review, critical analysis and editing were carried out by FB, TB. and EV. Funding acquisition was the responsibility of FB and TB. All authors read and approved the paper.

## ACKNOWLEDGMENTS

Drs Isabelle Léna and Massimo Mantegazza kindly provided Scn2a^+/-^ founder mice. The authors thank Drs Angelina Rogliardo and Antoine Besnard for insightful comments on the project, Dr Bruno Colombet for help with AnyWave software, Alicia Janvier for technical support and the iExplore animal facility at IGF, Montpellier. MGX acknowledges financial support from France Génomique National infrastructure, funded as part of “Investissement d’Avenir” program managed by Agence Nationale pour la Recherche (contract ANR-10-INBS-09). CR was funded by fellowships from the French Ministry for Higher Education and Research and from the French League Against Epilepsy (LFCE).

---

**Figure S1:**
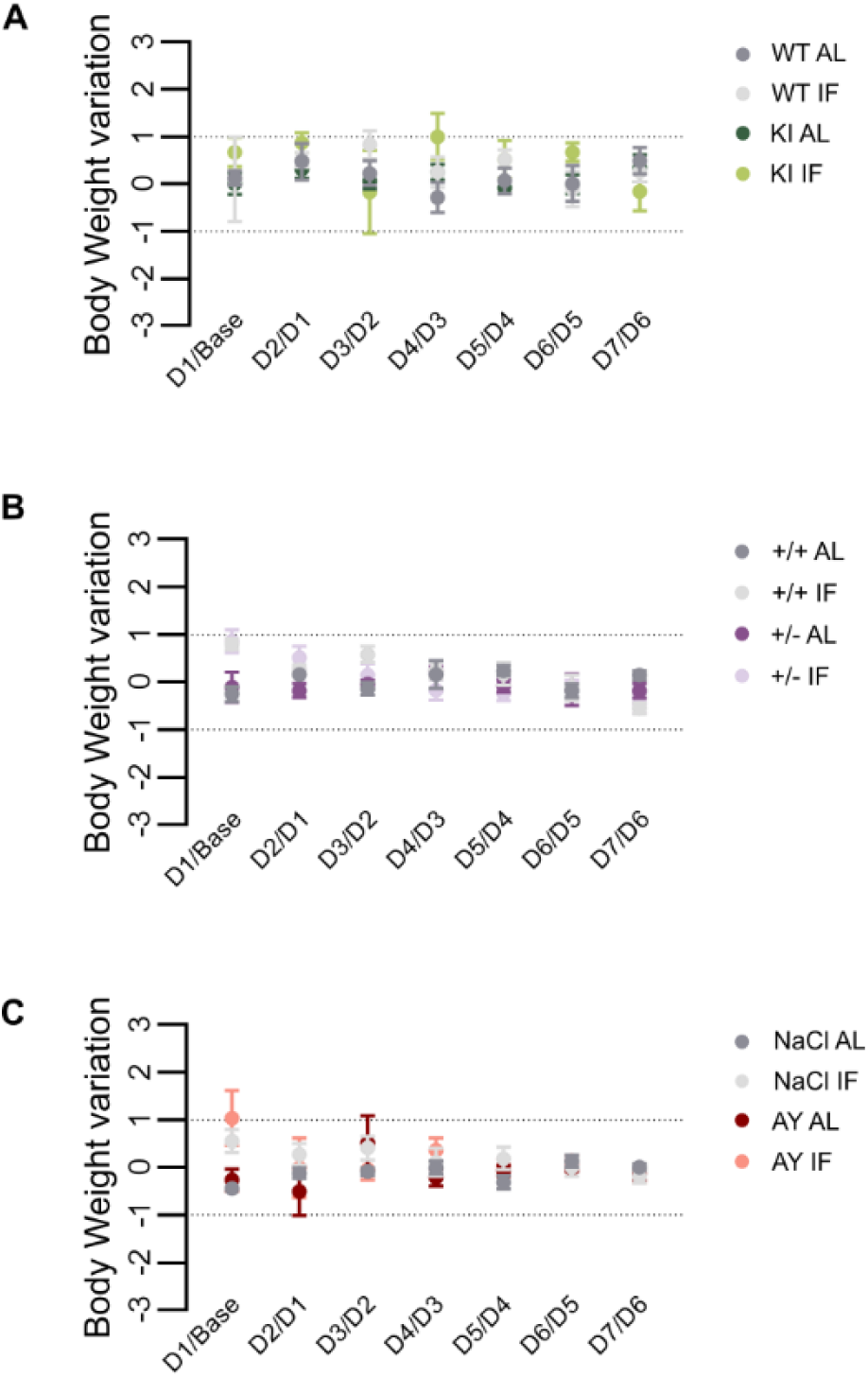
related to Figure 1: Body weight variation on week 1 of IF for Grm7^AAA^ WT and KI mice (A), Scn2A^+/+^ and ^+/-^ mice (B), AY-9944 mice and their NaCl controls (C).

**Figure S2:**
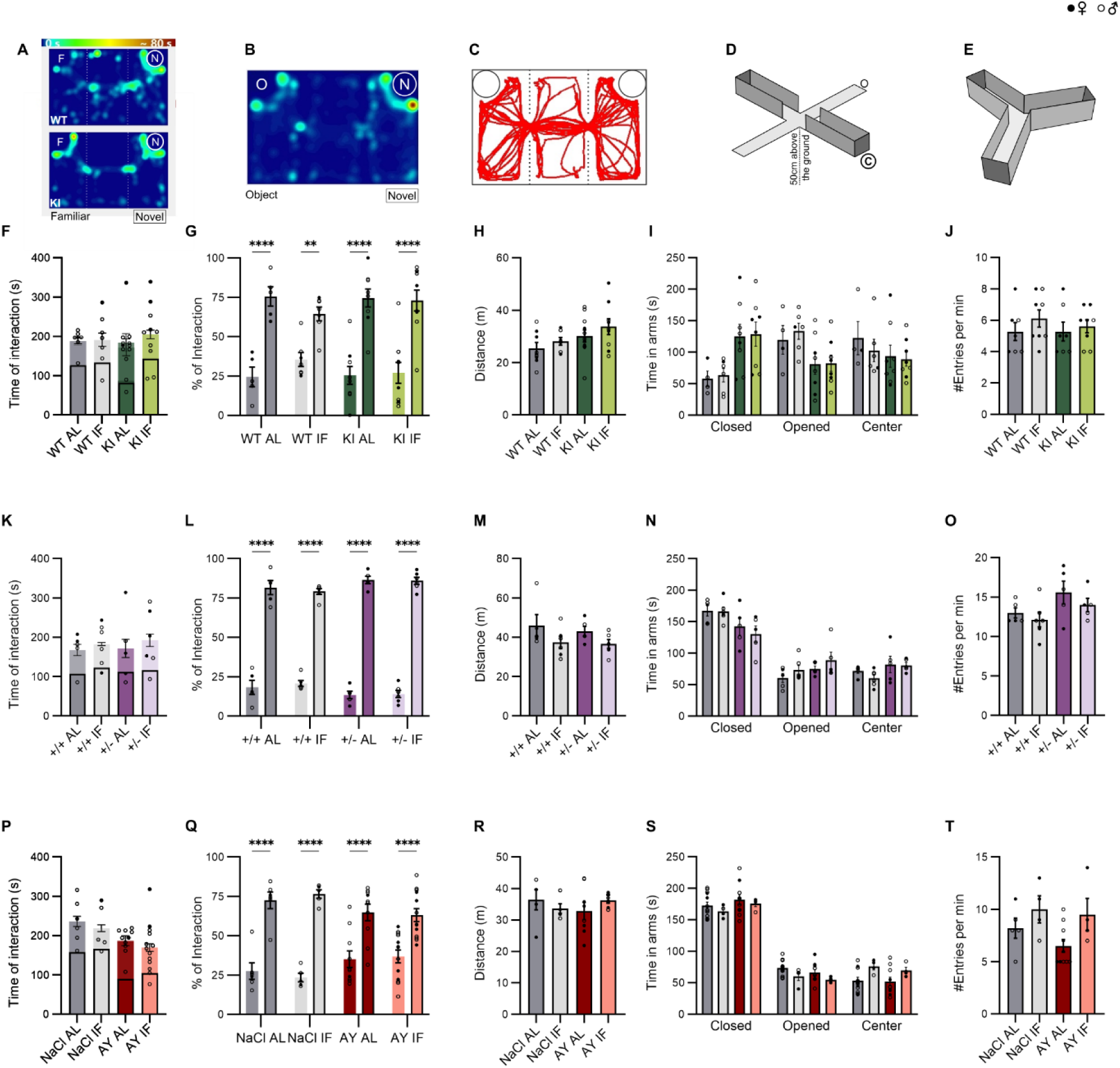
related to Figure 3: Additional behavioral tests parameters. A) Heatmap of the Crawley’s three-chamber social interaction test arena, during exploration by a WT mouse (top) and a KI mouse (bottom). B) Heatmap of the Crawley’s arena during the 2^nd^ phase of the trials. One cage contains an object, the other a novel mouse. C) Trace of a WT mouse in the Crawley’s arena during the habituation, when both cages are empty. D) Elevated plus maze apparatus with open (O) and closed (C) arms. E) Y maze apparatus. F) Total time of interaction with the novel and familiar stimuli for the Grm7 model, Scn2a model (K) and AY model (P). G, L, Q) Percentage of interaction with the object or novel mouse for each mouse strain and regime. H, M, R) Distance travelled in the empty Crawley’s arena for each mouse strain and regime. I, N, S) Total time spent in the center, closed and opened arms for each mouse strain and regime. J, O, T) Number of entries per minute in the arms of the Y-Maze for each mouse strain and regime. ** p<0.01, **** p<0.0001, Two-way ANOVA followed by Šidák test.

**Figure S3:**
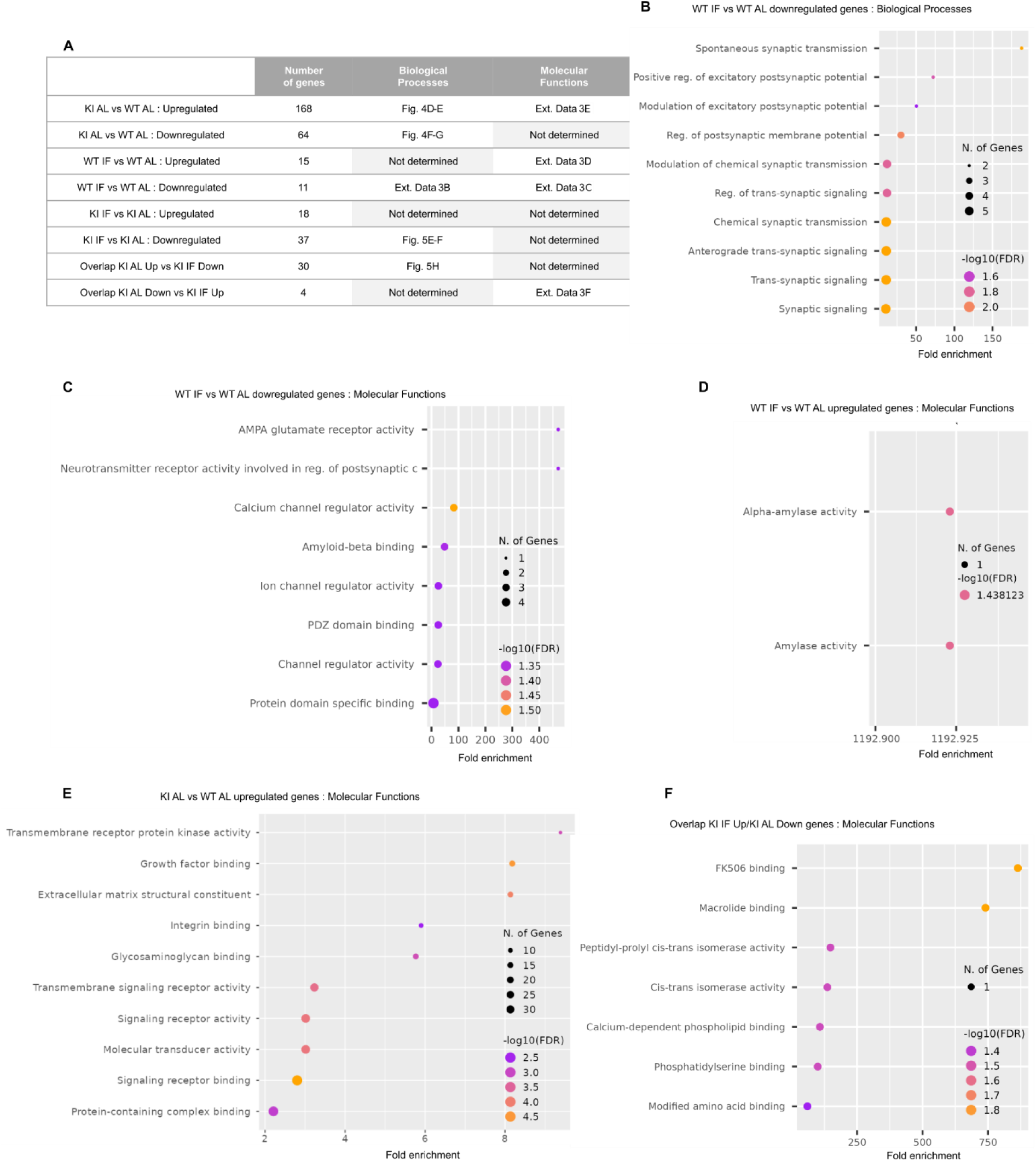
related to Figure 5: Additional Gene Ontology analysis of RNASeq data from Grm7^AAA^ WT and KI mice under AL or IF regime. A) Summary of the number of up- and downregulated DEG (DESeq, p<0.05) across conditions. The corresponding figure numbers where biological processes were assayed is indicated. NS: No specific biological process or molecular function was identified by the GO analysis. B, C) Biological processes (B) and molecular functions (C) associated with genes <u>downregulated</u> by IF in Grm7^AAA^ WT animals. D) Molecular functions associated with genes <u>upregulated</u> by IF in Grm7^AAA^ WT animals. E) Molecular functions associated with genes <u>upregulated</u> by IF in Grm7^AAA^ KI animals. F) Molecular functions enriched in overlapping genes downregulated in epileptic KI AL-fed animals and upregulated in KI IF-fed animals.

**Figure S4:**
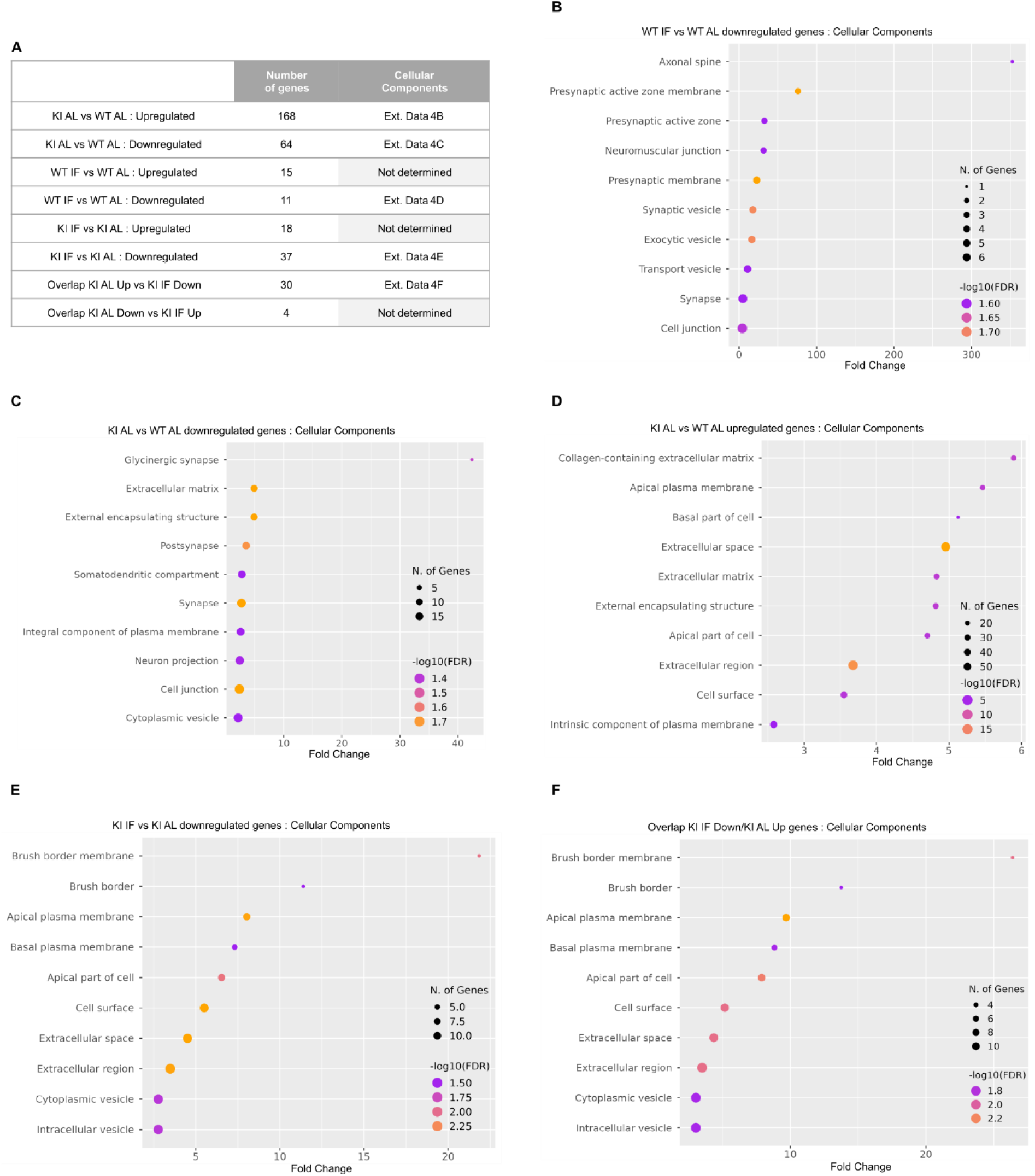
related to Figure 5: Cellular functions in Grm7^AAA^ WT and KI mice under AL or IF regime. A) Summary table of up- and down-regulated gene numbers across conditions, with the corresponding figure panel of GO analysis. NS: no specific cellular process was identified. B) Cellular processes associated with genes <u>downregulated</u> by IF in Grm7^AAA^ WT animals. C) Cellular processes associated with genes <u>upregulated</u> (C) and <u>downregulated</u> (D) by IF in Grm7^AAA^ KI epileptic animals compared to WT animals. E) Cellular processes associated with genes <u>downregulated</u> by IF in Grm7^AAA^ KI epileptic animals. F) Cellular functions identified for overlapping genes <u>upregulated</u> in epileptic KI AL-fed animals and <u>downregulated</u> in KI IF-fed animals.

**Figure S5:**
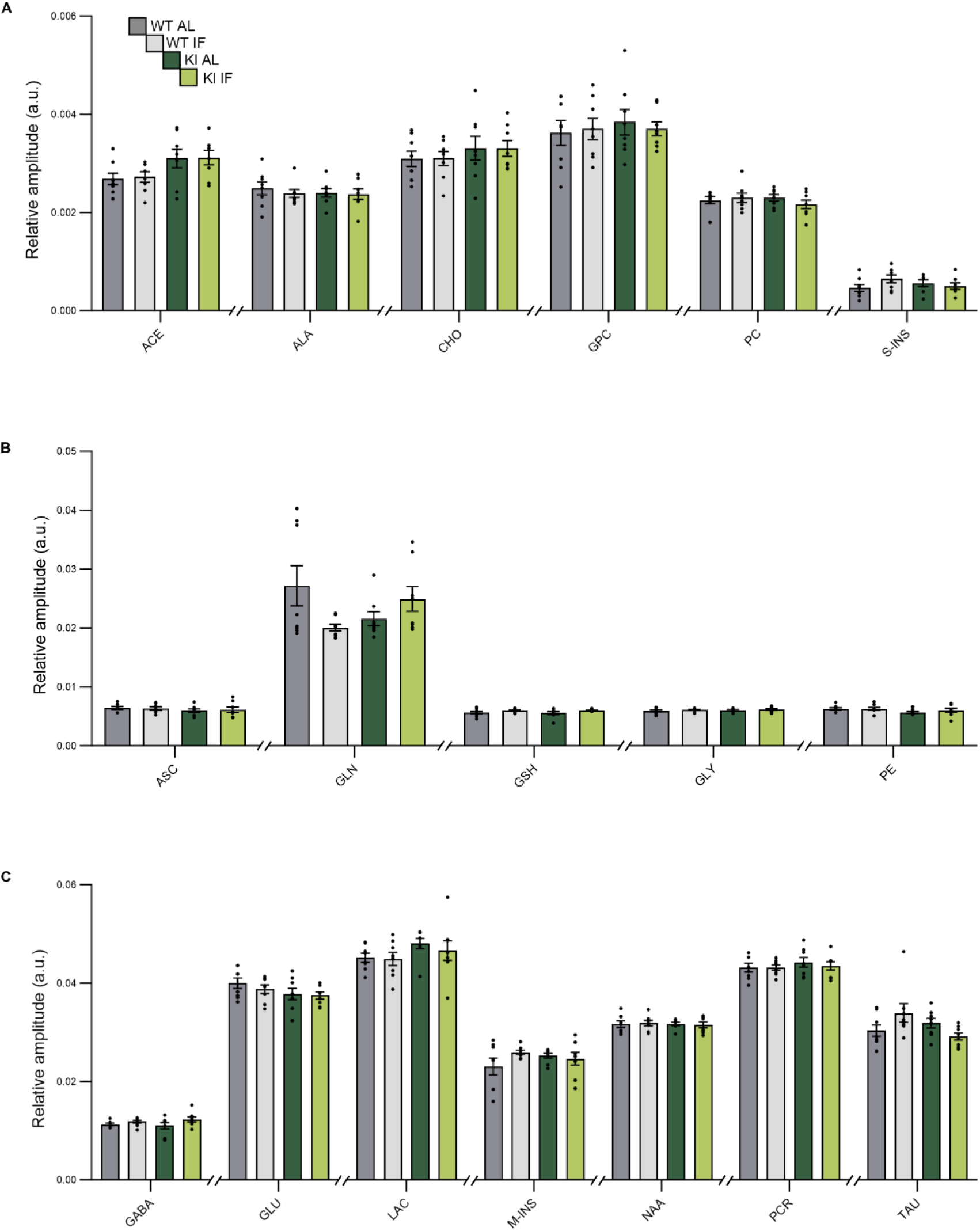
Metabolites detected in the thalami by *ex vivo* NMR from Grm7^AAA^ WT and KI mice under AL or IF regime. ACE acetate, ALA alanine, ASC ascorbate, CHO choline, CR creatine and phospho-creatine, GLU glutamate, GLN glutamine, GLY glycine, GPC glycerophosphocholine, GSH glutathion, LAC lactate, M-INS myo-inositol, NAA N-acetyl aspartate, PC phosphorylcholine, PE phosphoetaloamine, S-INS scyllo-inositol, TAU taurine. A two-way ANOVA followed by a Tukey post-hoc test for pair-wise comparison with correction for multiple comparisons was applied to evaluate the metabolic variations due to the regime in Grm7AAA KI animals compared to their WT controls.

**Supplemental Table 1:**
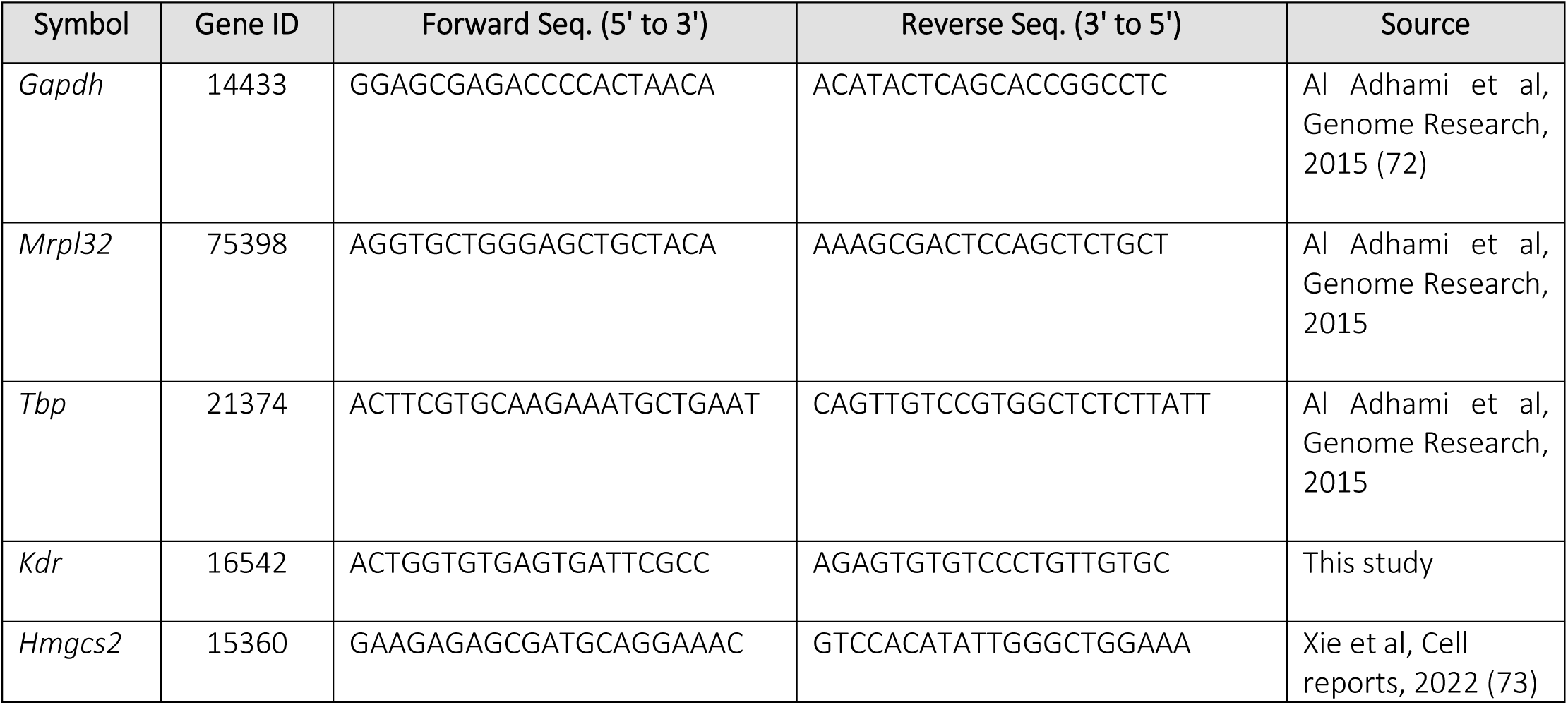

